# Neighborhood air pollution is negatively associated with neurocognitive maturation in early adolescence

**DOI:** 10.1101/2023.04.28.538763

**Authors:** Omid Kardan, Chacriya Sereeyothin, Kathryn E. Schertz, Mike Angstadt, Alexander S. Weigard, Marc G. Berman, Mary M. Heitzeg, Monica D. Rosenberg

**Affiliations:** University of Michigan, Department of Psychiatry, Ann Arbor, MI; University of Chicago, Department of Psychology, Chicago, IL; Chulalongkorn University, Faculty of Medicine, Bangkok, Thailand; University of Michigan, Department of Psychology, Ann Arbor, MI; University of Chicago, Neuroscience Institute, Chicago, IL

**Keywords:** Air pollution, Neurocognitive development, Functional brain connectivity, Cortical thickness, Early adolescence

## Abstract

Adolescence is a key period of neurocognitive maturation. Exposure to high levels of air pollutants have been associated with brain differences in youth, though the relevance of these brain findings to behavioral outcomes such as cognitive development is less clear. In this study, we used the US Environmental Protection Agency’s thresholds for unhealthy levels of fine particulate matter (PM_2.5_), Ozone (O_3_), and NO_2_ pollutants to compare youth exclusively exposed to high levels of each pollutant to their respective socioeconomically-matched low-pollution peers over a two-year period in the Adolescent Brain Cognitive Development (ABCD) Study. No youth in ABCD study were found to be at or above the unhealthy threshold for NO_2_. Separate multivariate analyses for PM_2.5_ (*N_High_*=348; *N_control_*=279) and O_3_ (*N_High_*=355; *N_control_*=324) resulted in two very similar neurocognitive latent variables loading positively on cortical functional maturation and task performance, and negatively on cortical grey matter thickness.

We found a significant difference in this neurocognitive maturation latent variable over time between the high-pollution and control groups from 9-10 to 11-12 years of age, such that maturation of cortical networks, increase in task performance, and cortical thinning were significantly higher in the control groups. These results were adjusted for parental income and education and youth’s age, sex, race/ethnicity, site, head-motion, scanner, general factor of psychopathology, pubertal status, and area deprivation index, in addition to the matching between high-pollution and control groups. In conclusion, exposure to high levels of PM_2.5_ and O_3_ is associated with lags in normative neurocognitive maturation in early adolescence.

## Introduction

The ability to maintain focus and process task-relevant information continues to develop during adolescence [Fortenbaugh et al., 2015; Isbell et al., 2015; Theodoraki et al., 2020; Ferguson et al., 2021], coinciding with functional and structural changes in the cortex [Dumontheil, 2016; Keller et al., 2022; Kardan et al., 2025]. Exposure to air pollutants as potential environmental neurotoxins has been investigated in multiple neuroimaging studies in this important period of neurocognitive development [see Morrel et al., 2025 for a review].

However, the specific link between these pollution-brain associations and cognitive development are unclear due to neuroimaging findings that are inconsistent across studies and sometimes at odds with well-known features of adolescent neurodevelopment [Parenteau et al., 2024; Sukumaran et al., 2025]. Additionally, the levels of pollution concentrations represented across studies vary widely, making them challenging to directly compare when results are inconsistent [Herting et al., 2019; 2024]. The goal of the current study is to address these gaps by utilizing theory-driven functional and structural measures of cortical maturation in conjunction with cognitive task performance to understand associations between high air pollution exposure, indicated by the U.S. Environmental Protection Agency thresholds, and neurocognitive maturation in youth.

In recent years, neuroimaging studies have started to characterize the association between fine particulate matter (PM2.5), surface Ozone (O3), nitrogen oxides (particularly NO2), and other air pollutants with both structural and functional brain measures in youth (see Herting et al., 2019; Morrel et al., 2025; and Parenteau et al., 2024 for reviews). In the structural domain, cortical grey matter thickness, white matter volume, as well as grey matter volume of subcortical regions may be associated with childhood exposure to PM2.5, though the direction of the associations are inconsistent across studies (e.g., compare Cserbik et al., 2020 and Pujol et al., 2016; compare Miller et al., 2022 and Beckwith et al., 2020; see Kusters et al., 2025 for no associations with white matter, grey matter, or subcortical volumes in a large sample from the Generation R study). In the largest study utilizing the Adolescent Brain Cognitive Development (ABCD^®^) Study baseline data, collected when youth were 9-10 years old, Sukumaran and colleagues reported associations between greater exposure to PM2.5 and NO2 and smaller surface area in the right inferior frontal gyrus but larger surface area in the right inferior frontal lobe [Sukumaran et al., 2025]. This study also found specific components of the PM2.5 to be differentially related to thinner or thicker cortical subregions [Sukumaran et al., 2025]. In addition to large sample size, strengths of this study include careful consideration of covariates and utilizing a multivariate approach. However, this study did not utilize the cognitive task or longitudinal data, making it difficult to assess neurodevelopmental trajectories (such as in cortical thinning) and neurocognitive changes.

In the functional brain connectivity domain, two studies using large samples have reported some inconsistent associations with PM2.5 and NO2 pollutants (Perez-Crespo et al., 2022; Cotter et al., 2023). Especially relevant to the current longitudinal study, Cotter and colleagues found associations between inter- and intra-network functional connectivity of specific networks (averaged for adult-defined Salience, Default mode, and Fronto-Partietal networks, as well as subcortical regions) and multiple pollutants (including PM2.5, O3, and NO2) using longitudinal ABCD Study data, collected when youth were 9-10 and 11-12 years old [Cotter et al., 2023]. Their findings were inconsistent with Perez-Crespo et al. [2022], which used cross-sectional fine-grained connectivity-level data (i.e., not averaged by network) from the Generation R study collected at ages 9-12. For example, higher segregation of (adult-defined) brain networks was associated with higher NO2 in Cotter et al. [2023], which is the opposite of the Perez-Crespo et al. [2022]’s findings. Additionally, Cotter and colleagues found associations between functional connectivity and PM2.5, while Perez-Crespo et al. [2022] did not.

Although some of these studies used careful covariate controls, they did not establish associations between pollution-linked brain network differences and cognitive performance or other behavioral measures. Additionally, Cotter et al. [2023] reported distinct functional connectivity changes based on exposure to PM2.5, O3, and NO2, with the findings not following the general hypothesis of neurodevelopmental delays being associated with higher pollution exposure. These may be due to their use of aggregated intra- and inter-network connectivity in Salience, Default mode, and Fronto-Partietal networks, even though boundaries of all of these networks are defined in adult samples. As the shifting of the early-life to adult network boundaries is still in progress in pre-to early adolescence [Kardan et al., 2025], averaging over adult-defined networks as units of analysis may reduce the signal of interest.

To address these gaps, namely lack of use of behavioral task data and using adult-defined networks when studying neurodevelopmental maturation in youth, we utilized longitudinal brain and cognitive task measures from the ABCD Study to understand the potential influence of exposure to high levels of PM2.5, NO2, and O3 within the US on normative neurocognitive maturation. We did not find any ABCD Study participants with NO2 exposure at or higher than the US Environmental Protection Agency (EPA)’s unhealthy threshold (though in general, surface O3 concentration is highly dependent on the availability of NOx in the air).

Therefore, NO2 was excluded from our analyses. Our hypothesis was that exposure to high levels of PM2.5 or O3 would be negatively associated with neurocognitive maturation among youth.

To test this hypothesis, we operationalized the normative cortical maturation process in both functional and structural domains. In the structural domain, cortical thinning is one of the most robust indicators of cortical maturation through adolescence [Squeglia et al., 2013; Fuhrmann et al., 2022; Bethlehem et al., 2022] and therefore the longitudinal cortical grey matter thickness values were used to assess the extent of cortical thinning between the high-pollution and control groups. In the functional domain, the distance from the canonical early-life neural connectivity profile and the proximity to adult brain network connectivity profile in the youth’s cortical functional connectome has recently been shown to capture neurocognitive maturation status both across and within youth [Kardan et al., 2025]. Therefore, this measure of cortical connectomic maturation was also longitudinally computed to assess the extent of normative cortical maturation between groups across timepoints. In addition to these neurodevelopmental measures, we also assessed cognitive performance in multiple tasks to capture the behavioral component of neurocognitive maturation, with the tasks longitudinally probing attention, working memory, and cognitive control development.

We employed an exposure vs. control approach to the data to allow for a more specific isolation of the pollution influences. Specifically, in larger multi-site studies, while the heterogeneity in the sample and higher variability in air pollution allow for more robust models, the confounding impacts of socioeconomic factors (SES) at the individual family and census tract level with air pollution introduce additional challenges that limit some of the conclusions that can be drawn. For example, if detrimental levels of a certain pollutant are nonexistent for certain SES combinations, then the relationship between a brain measure and pollution becomes artificially SES-dependent in the sample even if that is not true in the population. This would mean that using SES as a covariate may not remove all of its confounding effects due to the violation of the homogeneity of regression slopes assumption. The differential SES/Race/ethnicity composition of individuals residing near sources of particulate matter vs. those near sources of nitrogen oxides (and consequently O3) in the US additionally complicates efforts to isolate effects of air pollutants specifically and might drive some of the inconsistency between brain correlates of these pollutants. Therefore, we used a high-pollution vs. matched-control analysis approach to reduce the SES propensity difference for exposure to detrimental levels of each pollutant, while also including SES measures as covariates in the analyses.

Finally, we used specific thresholds of high “unhealthy” pollution provided by the EPA (annual mean > 9 µg/m^3^ for PM2.5; annual mean > 53 ppb for NO2; annual max > 70 ppm for O3; see Methods). Given the wide ranges of pollution levels across air quality neuroimaging studies, this approach makes comparing findings easier as it makes reproducing the boundary conditions of the current study possible for other researchers using data with partially overlapping pollution levels.

In sum, the goal of the current study is to isolate and understand the effects of exposure to high levels of fine particulate matter and ozone in pre- to early adolescence on normative cortical development and cognitive performance.

## Methods and Materials

### 1.1. Participants

We utilized data from the Adolescent Brain Cognitive Development (ABCD) Study^®^ [Casey et al., 2018], which is an ongoing longitudinal study of 11,867 children across 21 sites in the U.S. Participants were enrolled in the study between 9–10.9 years of age (Y0) and we additionally utilized their two-year follow-up (Y2; 11-12.9 years) data. Participants who passed fMRI functional connectivity and structural data quality checks and had cognitive task data (N-back and NIH-Toolbox; see Variables) at both baseline (N = 7396) and follow-up (N = 5170) for our longitudinal analyses were a total of N = 3645. These provided the pool of participants that were used across studies 1 and 2. These participants were comparable to the total sample in terms of air pollution exposure, and included 15.8% exposed to high PM2.5 (compared to 16.7% in the total sample), 19.1% exposed to high O3 (compared to 12.1% in the total sample), and 0% exposed to high NO2 (compared to 0% in the total sample). Since the ABCD Study sample is from the US, we used the EPA thresholds. Whether the EPA’s standard for unhealthy NO2 concentration is set too high or not is outside of the scope of the current study, but using the World Health Organization (WHO) threshold would result in almost all of the sample being considered above the recommended level (see thresholds in the Variables section; see [Herting et al, 2024] for comparing U.S. EPA and the WHO pollution thresholds).

To isolate the PM2.5 and O3 effects from each other, participants who had both high PM2.5 and high O3 exposures were also removed from the pool of participants (N = 40). Additionally, 35 high-PM2.5 participants were excluded because they had no same-demographic peers among the lower pollution participants (mean PM2.5 = 9.70 µg/m3 in the excluded high-PM2.5 participants compared to 9.70 µg/m3 in the included high-PM2.5 participants). The first study included 348 participants with High-PM2.5 exposure and 279 participants in the control group.

The latter were randomly selected from the pool of low-PM2.5 participants, while being matched with the High-PM2.5 group in terms of household income, parental education, race/ethnicity, biological sex, and age. Full matching was not possible without introducing duplicates in the control group, hence the final N for them is 279 rather than 348. The matching successfully reduced the differences between high-PM2.5 group and low-PM2.5 pool of participants such that the matched control group’s differences in all domains had Cohen’s ds ≤ 0.2 (see top half of **Table 1** for demographic characteristics of the high-PM2.5 group and its comparison to the matched group and the pool of low-PM2.5 participants). The small remaining differences were not possible to remove via matching, but matching was supplemented by having these variables, among others, as covariates in the multivariate analyses.

**Table 1.** Demographic break-down of the participants in Study 1 (top half; PM2.5) and Study 2 (bottom half; O3). Notes: Mean household income was approximated by interpolating mean of the midpoints of income brackets. Min and max income are shown in brackets. Education is based on the highest parental years of education. Bold values show Cohen’s d > 0.2 prior to matching.

|  | High-PM2.5 group<br>(N = 348) | Control low-PM2.5 group matched with High-PM2.5<br>(N = 279) | Low-PM2.5 pool<br>(N = 1381) | Cohen's d<br>(high-PM2.5 vs. control) | Cohen's d<br>(high-PM2.5 vs. low-PM2.5 pool) |
| --- | --- | --- | --- | --- | --- |
| **Parental Education** | 16.8 yrs<br>(SD = 2.4) | 17.2 yrs<br>(SD = 2.2) | 17.9 yrs<br>(SD = 2.0) | -0.17 | **-0.53** |
| **Household Income** | \$64.6K [5K-200K] | \$74.5K [5K-200K] | \$106K [5K-200K] | -0.20 | **-0.70** |
| **White** | 150 (43.1%) | 145 (51.9%) | 1000 (72.4%) | -0.18 | **-0.67** |
| **Hispanic** | 91 (26.1%) | 64 (22.9%) | 178 (12.9%) | 0.07 | **0.37** |
| **Black** | 50 (14.4%) | 26 (9.3%) | 77 (5.6%) | 0.15 | **0.42** |
| Asian | 10 (2.9%) | 6 (2.2%) | 14 (1.0%) | 0.05 | 0.14 |
| Other | 13.5% | 13.6% | 8.1% | 0.00 | 0.19 |
| Male | 174 (50.0%) | 143 (51.2%) | 710 (51.4%) | -0.03 | -0.04 |
| Female | 174 (50.0%) | 136 (48.8%) | 671 (48.6%) | 0.03 | 0.04 |
| Age at Y0 | 9.92 yrs<br>(SD = 0.65) | 9.97 yrs<br>(SD = 0.63) | 10.00 yrs<br>(SD = 0.61) | -0.09 | -0.14 |
| Age at Y2 | 11.94 yrs<br>(SD = 0.69) | 12.04 yrs<br>(SD = 0.68) | 12.07 yrs<br>(SD = 0.64) | -0.16 | -0.19 |
|  | High-O3 group<br>(N = 355) | Control low-O3 group matched with High-O3<br>(N = 324) | Low-O3 pool<br>(N = 978) | Cohen's d<br>(high-O3 vs. control) | Cohen's d<br>(high-O3 vs. low-O3 pool) |
| Parental Education | 17.8 yrs<br>(SD = 2.2) | 17.9 yrs<br>(SD = 2.1) | 17.5 yrs<br>(SD = 2.4) | -0.06 | 0.12 |
| Household Income | \$102.5K [5K-200K] | \$104.8K [5K-200K] | \$90.4K [5K-200K] | -0.02 | 0.11 |
| White | 254 (71.5%) | 243 (75.0%) | 633 (64.7%) | -0.08 | 0.09 |
| Hispanic | 16% | 14.2% | 17.6% | -0.05 | 0.05 |
| Black | 1.9% | 1.9% | 8.2% | 0.00 | -0.18 |
| Asian | 1% | 1% | 2.0% | 0.02 | -0.10 |
| Other | 9.3% | 8.0% | 7.5% | 0.05 | 0.12 |
| **Male** | 55.2% | 52.5% | 47.3% | 0.05 | **0.22** |
| **Female** | 44.8% | 47.5% | 52.7% | -0.05 | **-0.22** |
| Age at Y0 | 9.99 yrs<br>(SD = 0.63) | 9.97 yrs<br>(SD = 0.62) | 9.97 yrs<br>(SD = 0.61) | 0.04 | 0.04 |
| Age at Y2 | 12.06 yrs<br>(SD = 0.63) | 12.03 yrs<br>(SD = 0.65) | 12.02 yrs<br>(SD = 0.64) | 0.04 | 0.06 |

The second study included 355 participants with High-O3 exposure and 324 with low O3 exposure who were randomly selected from the low-O3 pool and matched with the High-O3 group (see bottom half of **Table 1** for demographic characteristics of these groups). In general, unlike high vs. low PM2.5 which had large differences in SES and race/ethnicity, the high vs. low O3 participants were not as different in these dimensions and the matching only balanced the group sizes (and matched the small difference on biological sex).

### 1.2. Resting-state fMRI processing pipeline and exclusions

For both Y0 and Y2 neuroimaging sessions, resting-state fMRI was acquired in four runs (∼5 min per run, full details are described in [Hagler et al., 2019]). Details of the pre-processing pipeline can be found in previous work [Sripada et al., 2021]. Key features of the pipeline include FreeSurfer normalization, ICA-AROMA denoising, CompCor correction, and censoring of high motion frames with a 0.5 mm framewise displacement threshold. Visual quality control (QC) was conducted to assess registration and normalization steps. Participants with at least one resting-state run in each year (Y0 and Y2) with each run having more than 250 degrees of freedom left in the BOLD timeseries after confound regression and censoring were included in the analyses. The resting-state fMRI data were spatially down-sampled to 333 parcels in the Gordon parcellation for the cortical regions [Gordon et al., 2016] and cortical functional connectivity matrices were generated for each participant for baseline and Y2 by Fisher Z transformation of the 333*333 correlation matrices.

### 1.3. Variables

#### Cortical functional connectomic maturation

Anchored functional connectivity maturation (AFC) score was calculated for the resting-state functional connectivity matrix of each participant at each year (Y0 and Y2). AFC captures cortical functional connectomic maturation in youth by arranging the connectivity matrix according to both early-life set of networks and adult set of networks and comparing the distance of the connectome to these two anchors [Kardan et al., 2025]. This measure considers the boundaries of brain functional networks during adolescence in transition from early-life to adulthood and, due to its two anchoring points, is significantly better than distance from adult networks alone in capturing age and ageing in youth [Kardan et al., 2025]. Importantly, individual differences in AFC values across participants, as well as the amount of developmental change in AFC within a participant, are respectively positively associated with individual differences in cognitive performance and improvements in cognitive performance in youth [Kardan et al., 2025].

We used the adult [Gordon et al., (2016)] and baby [Kardan et al., (2022)] sets of networks as anchors following [Kardan et al., 2025]. These sets of networks and their anatomical maps are shown in **Figure 1** (above diagonals). These sets of networks both capture the youth connectomes at 9-13 years of age better than null models based on proximity or rotations (see [Kardan et al., 2025] for details). The maturity of the participant’s cortical networks is then calculated as their connectome’s proximity to adult and distance from baby networks.

**Figure 1.**
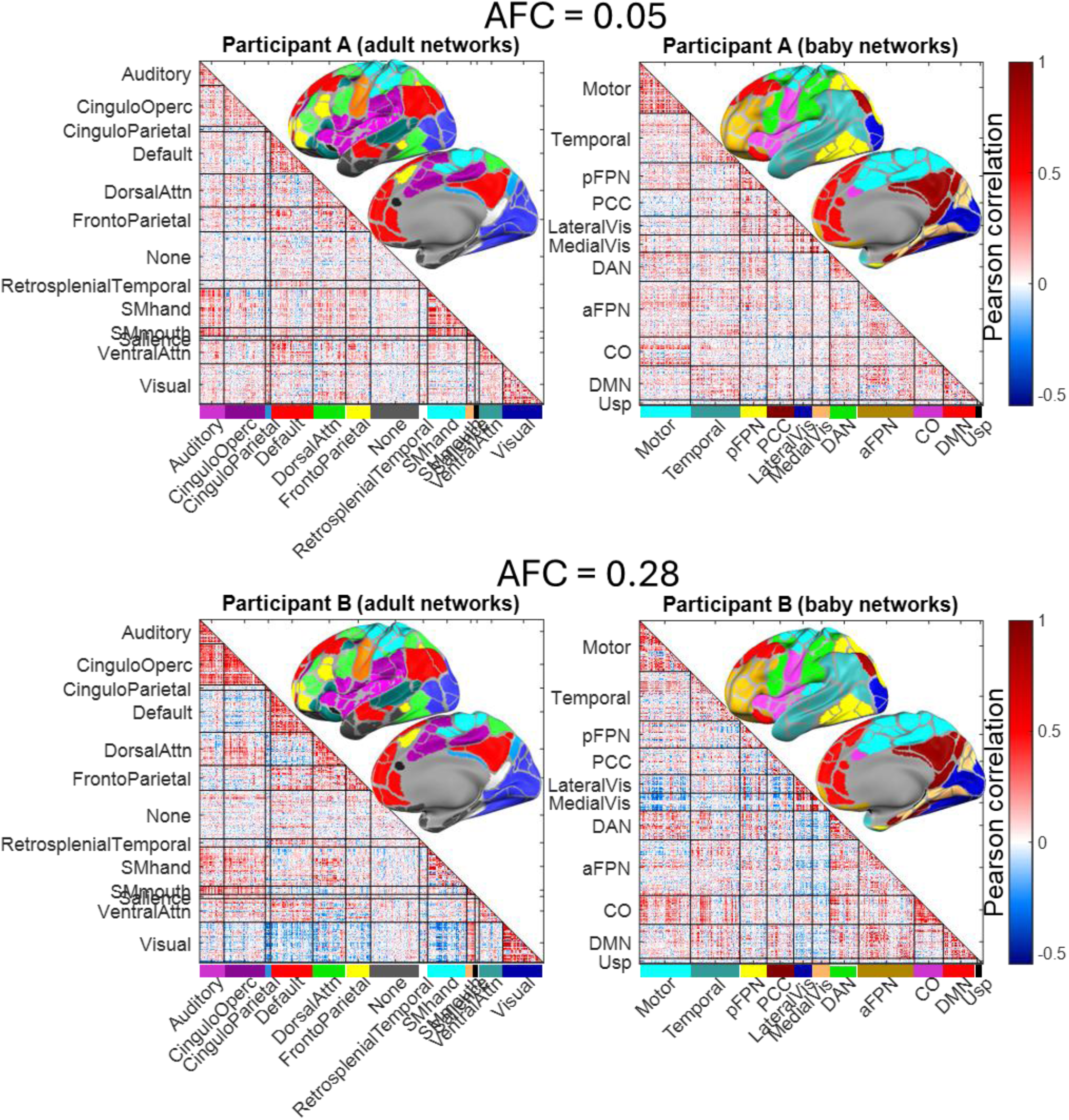
Above diagonals: Anatomical map of the Gordon, et al. 2016 (adult) and Kardan et al., 2022 (baby) networks. **Below diagonals:** Demonstration of Anchored functional connectivity (AFC) maturation score for two example participants at baseline. Participant B (AFC = 0.28) has moved further along the normative functional connectomic maturation in their cortex compared to Participant A (AFC = 0.05). Some values from within rectangular (between-network) and triangular (within-network) patches in the figures have been randomly shuffled to prevent unauthorized reconstruction of individual level data from the figure.

Specifically, AFC was calculated as follows for each participant at Y0 as well as at Y2:

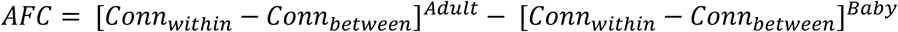

Above, [Conn_within –_ Conn_between_]^Adult^ is the average connectivity within networks minus the average connectivity between networks specified by [Gordon et al., 2016], while [Conn_within –_ Conn_between_]^Baby^ is the average connectivity within networks minus the average connectivity between networks specified by [Kardan et al., 2022].

Figure 1 compares two example participants’ connectivity matrices at the same age and at baseline (participants A and B). Connectomes are arranged according to the adult (left) and baby (right) networks. The better aggregation of higher values (red) within adult networks (triangular patches in the connectivity matrices) compared to within baby networks results in a large positive AFC value for participant B. On the other hand, the relatively similar aggregation of higher values within adult or baby networks in participant A results in a close-to-zero AFC for participant A (Negative AFC would mean baby networks fit the connectome better than adult networks). Figure 2 shows the distributions of AFC values across ABCD study participants at both Y0 and Y2 (these are previously shown in Kardan et al., 2025 and are reported here for the longitudinal sample included in the current study).

**Figure 2.**
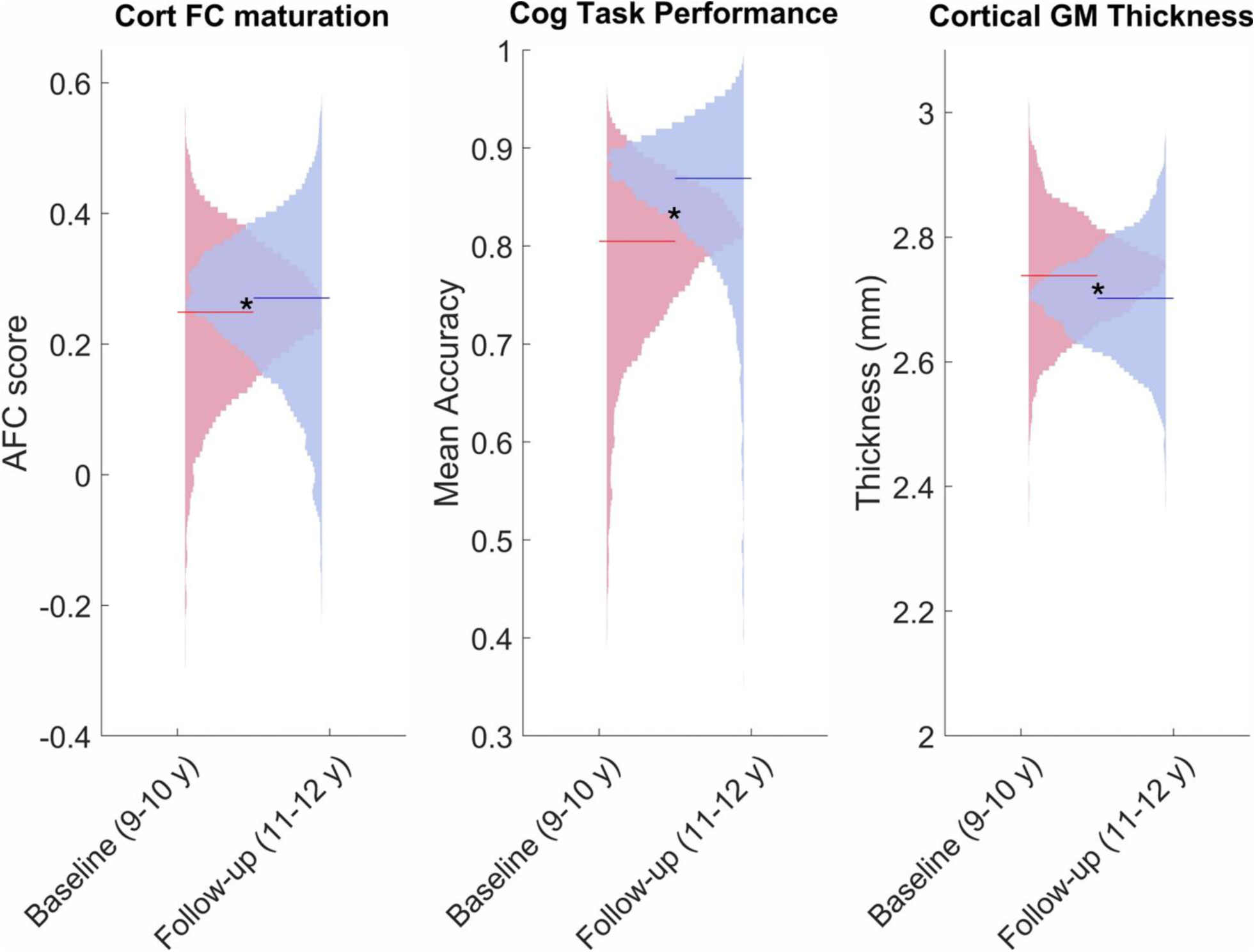
The distributions of the three variables used to assess neurocognitive maturation across the entire longitudinal sample of the current study (N = 3645). Consistent with previous work using ABCD Study sample, cortical functional connectomic maturation and task performance both significantly increased over the two-year period as expected. Cortical thickness decreased across the two years as expected. Note: * indicates p < 0.001 based on paired t-test (N = 3645).

#### Cortical thickness

The average cortical thickness for Y0 and Y2 of each participant was available based on the generated variable ‘smri_thick_cdk_mean’ by the ABCD Data Analysis, Informatics & Resource Center (DAIRC), which is the average thickness in mm over all the cortical parcels of both hemispheres based on the Desikan-Killiany atlas [Desikan et al., 2006]. Mean cortical thickness peaks as early as 2 years of age and normatively declines in close to linear fashion during development [Bethlehem et al., 2022], hence lower values at Y2 compared to Y0 are expected to reflect cortical structural maturation. The mean cortical thickness values across all included participants are shown in Figure 2.

#### Cognitive task performance

Cognitive performance was quantified as average performance in the N-back task [Casey et al., 2018], and five of the NIH-Toolbox [Gershan et al., 2013] tasks: the Picture Vocabulary task, Flanker inhibitory control and attention task, Pattern Comparison processing speed task, Picture sequence memory task, and Oral reading recognition task. The other two of the NIH-Toolbox tasks (List Sorting and Dimensional Change Card Sort) were not collected at Y2 by the ABCD Study and therefore were not included. At each year, the uncorrected scores across NIH-Toolbox tasks were averaged and scaled to 0-1 performance and then averaged with the N-back accuracy (mean of 0-back and 2-back accuracies). These tasks were chosen because they capture a wide range of developing cognitive processes in youth including sustained attention, working memory, and inhibitory control and are collected in both ABCD data releases (Y0 and Y2). Using a Principal Component Analysis (PCA) to combine the N-back and NIH-Toolbox performances results in a first PC that is highly correlated with the mean of the tasks (r = 0.68 at Y0 and r = 0.69 at Y2), so the simpler method (i.e., averaging) which does not introduce data leakage was used. The mean task performance values across all included participants are shown in Figure 2.

#### Air pollution

The fine particulate matter (PM2.5), O3, and NO2 air pollution were from the led_l_pm25, led_l_o3, and led_l_no2 tables from the ABCD geo-data workgroup at baseline, respectively (annual average concentration at 1x1 Km^2^ around the primary residence for PM2.5 and NO2; maximum annual concentration for O3). Based on the EPA’s guidelines (https://www.epa.gov/criteria-air-pollutants/naaqs-table) for unhealthy levels of air pollutants, the thresholds of >9 µg/m^3^ for annual average PM2.5, >70 ppb for annual fourth-highest daily maximum 8-hour concentration for O3, and >53 ppb for annual average NO2 were chosen. We may be overestimating the high-O3 exposure since the ABCD Study variable for O3 is for the maximum (rather than 4^th^ highest) concentration over the year. Additionally, we found no ABCD Study participants at or exceeding the 53 ppb threshold for NO2, so only PM2.5 and O3 were included in this study. The low-pollution threshold was set at <7 µg/m^3^ for PM2.5 and <60 ppb for O3, respectively. These values were chosen based on incrementing the low-pollution thresholds during matching until they resulted in a large enough pool of participants to draw matched controls for high-PM2.5 and high-O3 groups (i.e., with Cohen’s |d| ≤ 0.2 across matching categories), respectively. The size of these pools of participants are reported in **Table 1**.

#### Covariates

In addition to sex, age, parental education, household income, and race/ethnicity, other covariates that the multivariate results were adjusted for included data collection site, head motion (mean framewise displacement), scanner type, number of fMRI runs, pubertal status, general factor of psychopathology, and area deprivation index. Pubertal status was the average of youth and parental reports on the Pubertal Developmental Scale categorized sum score; 1: pre-puberty, 2: early puberty, 3: mid puberty, 4: late puberty, and 5: post puberty. The general factor of psychopathology was from a bi-factor model fit to eight parent-rated Child Behavior Checklist age 6 to 18 form [Achenbach et al., 2000], (see factor model details in [Clark et al., 2021]). The area deprivation index [Kind and Buckingham, 2019] for each participant’s address was obtained from the weighted sum score linked by the ABCD Study geo-data workgroup and derived from the American Community Survey at census tract resolution to quantify neighborhood socioeconomic disadvantage.

### 1.4. Statistical analysis

#### Partial Least Squares (PLS) analysis

In the PM2.5 study, mean-centered task partial least squares (PLS) multivariate analysis [McIntosh et al., 2004; Krishnan et al., 2011; Kardan, Weigard, et al., 2025] was used to identify the combination of three neurocognitive variables (AFC, cortical thickness, and task performance) that are maximally related to the group (high-PM2.5 vs. matched control) by time (Y0 and Y2). PLS is computed via singular value decomposition applied to the covariance between the neurocognitive variables and the group-by-time contrasts. Applying mean-centering to either groups or timepoints can emphasize the temporal contrast versus the group differences, respectively. In the current analysis, the goal was to allow for a data-driven group-by-time contrast to emerge (i.e., allowing for potential group-by-time interaction). Therefore, only the grand mean was removed from the neurocognitive values prior to calculation of the covariance matrix X’Y, (where X represents the neurocognitive variables and Y represents the group-by-time contrasts). Applying SVD to the covariance matrix results in three latent variables (LVs) that are linear combinations of AFC, cortical thickness, and task performance that are differentially instantiated for different group-by-time cells.

To test the significance of each LV, 1000 covariance matrices were generated by randomly permuting condition labels for the X (neurocognitive) variables. These covariance matrices reflect the null hypothesis that there is no relationship between X and Y variables. They were subjected to SVD as before resulting in a null distribution of singular values. The significance of the original LV was assessed with respect to this null distribution. The weighted scores for the LV that passed the permutation test were correlated with the original neurocognitive values and group-by-time dummy variables, adjusted for sex, age, parental education, household income, race/ethnicity, data collection site, scanner type, number of fMRI runs, pubertal status, general factor of psychopathology, and area deprivation index. This turned the weights into more interpretable loadings (i.e., partial correlations) while also adjusting them for all the covariates. This was done 5000 times with bootstrapped samples to generate 95% confidence intervals for the adjusted PLS loadings. We only interpreted the primary LV, because the 2^nd^ LV did not have any significant loadings (i.e., bootstrap distributions contained zero for all partial correlations) and the 3^rd^ LV did not pass the permutation test (p > 0.05).

In the second study, the same PLS analysis was repeated for the high-O3 vs. its respective matched control group. Here again, only the primary latent variable passed both permutation and partial correlation bootstrap tests, and thus the other LVs are not discussed further.

## Results

### Fine particulate matter and neurocognitive maturation

The primary latent variable (LV) from the PLS analysis to differentiate the high-PM2.5 and its low-PM2.5 matched control group across the Y0 and Y2 is shown in Figure 3. The LV’s adjusted correlations (see Covariates in the Methods) with anchored functional connectivity maturation (AFC), cognitive task performance, and cortical thickness suggest that it reflects normative neurocognitive maturation in all three domains (Figure 3 left panel). Adjusted for all covariates, this neurocognitive maturation LV loaded significantly more on Y2 compared to Y0 within the low-PM2.5 control group (p_bootstrap_ = 0.015, N = 279), but not within the high-PM2.5 group (p_bootstrap_ = 0.820, N. S., N = 348) (see Figure 3 right panel). This interaction was significant (p_bootstrap_ = 0.001), such that the increase in AFC and task performance and decrease in cortical thinning within the control group was significantly more than that of the high-PM2.5 group (which did not show any significant difference between Y0 and Y2). The effect size for this increase in the neurocognitive LV within the two-year period was small within the control group as indicated by the change in partial correlation (Δr_partial_ = 0.114; p_bootstrap_ = 0.015).

**Figure 3.**
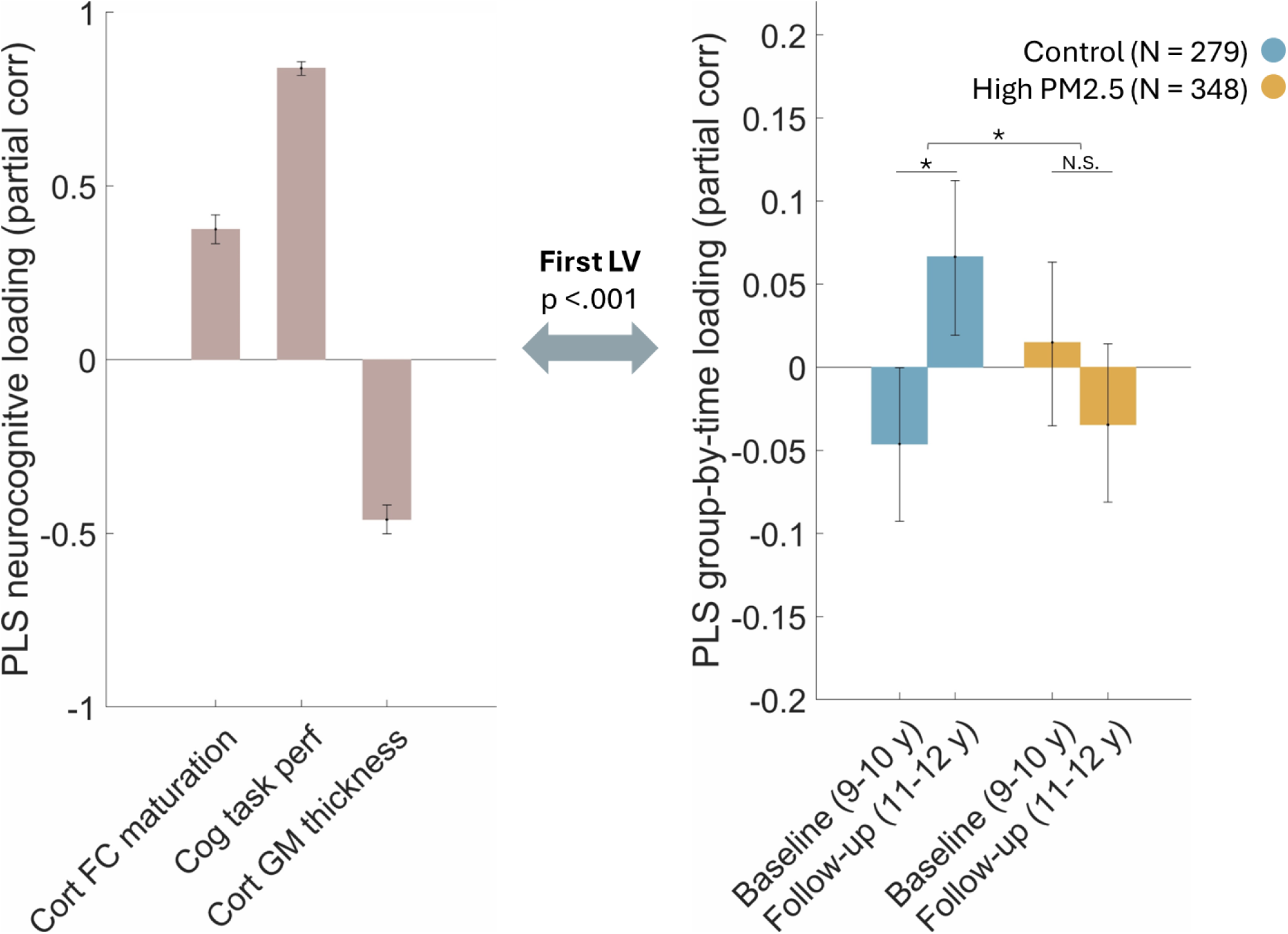
The primary latent variable from the PLS analysis differentiating the high-PM2.5 and matched control groups over the two timepoints (Y0:9-10 years and Y2:11-12 years) in terms of their neurocognitive measures. The primary LV was the dominant LV in the PLS with 86% of the covariance explained determined by σ_XY_ = 0.86 calculated as the squared singular value for this LV divided by the sum of squared singular values for all LVs. This is not a measure of effect size, and the partial correlations with group-by-time can be interpreted as effect sizes). The y-axes show partial correlation of the LV (adjusted for all covariates) with the neurocognitive measure (left panel) or group-by-time contrasts (right panel). Error bars show 95% confidence intervals based on 5000 bootstraps.

### Surface Ozone and neurocognitive maturation

The primary LV differentiating the high-O3 and its low-O3 matched control group across the Y0 and Y2 is shown in Figure 4. The LV’s adjusted correlations AFC, cognitive task performance, and cortical thickness again showed the LV as being consistent with normative neurocognitive maturation (Figure 4 left panel), similar to the LV for the PM2.5 analysis. Echoing the isolated PM2.5 results, adjusted for all covariates, this neurocognitive maturation LV loaded significantly more on Y2 compared to Y0 within the low-O3 control group (p_bootstrap_ = 0.021, N = 324), but not within the high-O3 group (p_bootstrap_ = 0.727, N. S., N = 355) (see Figure 4 right panel). This interaction was significant (p_bootstrap_ = 0.005), such that the increase in AFC and task performance and decrease in cortical thinning within the control group was significantly more than that of the high-PM2.5 group (which did not show any significant difference between Y0 and Y2). Here, again, the change in partial correlation (Δr_partial_ = 0.091; p_bootstrap_ = 0.021) indicated a small effect size for the increase in the neurocognitive LV within the two-year period for the low pollution control group.

**Figure 4.**
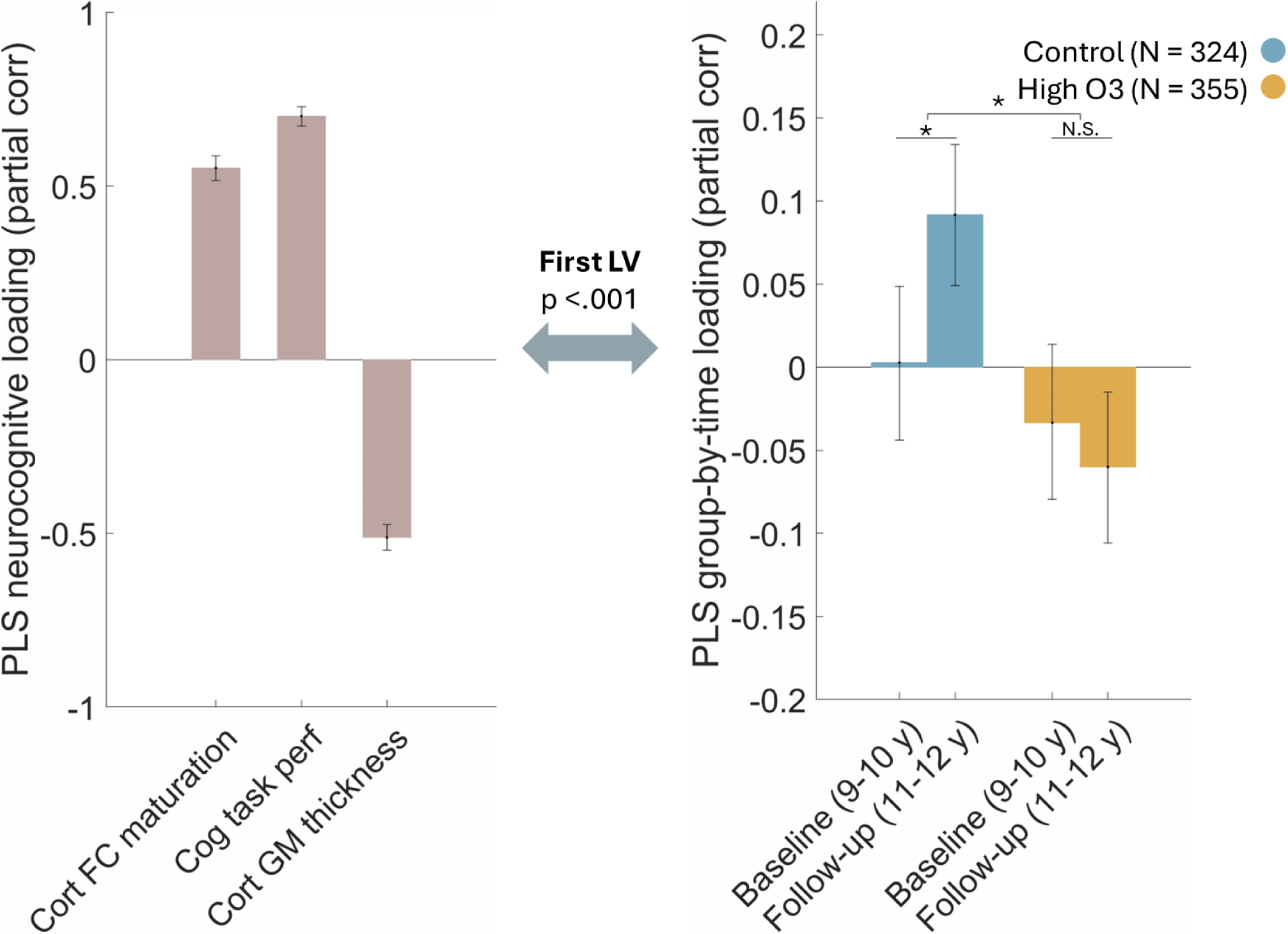
The primary latent variable from the PLS analysis differentiating the high-O3 and its matched control groups over the two timepoints (Y0:9-10 years and Y2:11-12 years) in terms of their neurocognitive measures. The primary LV was the dominant LV in the PLS with 94% of the covariance explained (This is not a measure of effect size, and the partial correlations with group-by-time can be interpreted as effect sizes). The y-axes show partial correlation of the LV (adjusted for all covariates) with the neurocognitive measure (left panel) or group-by-time contrasts (right panel). Error bars show 95% confidence intervals based on 5000 bootstraps.

## Discussion

In the current study, we utilized matched subsamples from a large and heterogenous longitudinal dataset (ABCD Study) to compare the neurocognitive development of youth exposed to high levels of fine particulate matter (annual mean PM2.5 > 9 µg/m^3^) or surface ozone (annual max day O3 > 70 ppb) compared to their low-pollution peers. In two separate analyses, we found small but robust effects showing significantly more neurocognitive maturation occurring from 9-10.9 years to 11-12.9 years of age within the low-pollution groups. These results were adjusted for a comprehensive list of covariates including family income, parental education, age, sex, race/ethnicity, site, pubertal status, general psychopathology, and area deprivation index in addition to the matching of control groups to the high pollution groups on the SES factors. The emerged neurocognitive latent variable was robust across analyses and loaded on structural (grey matter thinning), functional (cortical functional connections organizing further from early-life and closer to adult functional networks), and behavioral (improvements in cognitive task performance) domains of youth neurocognitive development.

In the structural MRI domain, multiple air pollution neuroimaging studies in recent years have investigated the longitudinal changes in gray and white matter development during early adolescence, with the directionality of associations being mixed (see [Morrel et al., 2025] for a review). Some of these inconsistencies have been attributed to timing of pollutant exposures and the specific types of pollutants (though qualitatively not meta-analytically). Inconsistent findings can additionally be as a result of variabilities across studies in how well the pollution associations were isolated from other confounding factors and the degree of SES propensity mismatch across levels of pollution in the sample. Using structural data at voxel-level granularity despite insufficient sample size could be another contributing factor to inconsistencies across some of these studies.

In the functional connectivity domain, however, our finding of lower AFC change in high-PM2.5 compared to control groups does generally follow some of the reported findings from [Zundel et al., 2024] and [Cotter et al., 2023]. Importantly, these studies—and all other studies investigating within- and between-network connectivity as a function of pollution—assume that adult networks capture the functional organization of 9-12-year-olds’ brains. Prior work demonstrates that this assumption may miss important aspects of the functional maturation process. Rather, functional connectome maturation in this age range is better measured as the distance of an individual’s connectome to a young adult vs. an infant network organization template [Kardan et al., 2025]. Therefore, to relate [Zundel et al., 2024] and [Cotter et al., 2023] findings to AFC one must pay attention to the network’s boundaries in early-life vs. adulthood (see Figure 1, above diagonals). In doing so, some of the reported findings associated with more PM2.5 in these studies can be qualitatively reinterpreted. For example, [Zundel et al., 2024] reports attenuated decrease in connectivity between adult Cingulo-opercular network and adult Ventral attention network associated with PM2.5. When considering the early-life networks, this can be re-interpreted as lack of expected segregation in the early-life Temporal network. They also report an increased connectivity between adult-DMN and adult-DAN associated with PM2.5, which seems to be lack of integration of the anterior and posterior components of DMN (labeled as PCC and DMN in early-life networks). Both of these would be considered delayed maturation and result in a lowered anchored FC maturation (AFC) score, which is in line with our PM2.5 analysis results. There are, however, other findings in [Zundel et al., 2024] that do not follow this neoteny-maturity axis used in AFC over the two-year period.

A key strength of our study is its utilization of longitudinal summary measures of the degree to which individuals show expected patterns of global cortical development. This keeps our neural measures interpretable and grounded in neurodevelopmental theory. Specifically, both shifting cortical functional organization from early-life towards adult network boundaries (increase in AFC) [Kardan et al., 2025] and cortical thinning due to myelination and synaptic pruning [Bethlehem et al., 2022] are expected to occur in the age range of the study. The alignment of the data-driven PLS latent variable with the theoretical expectation within the low-pollution control groups indicates that we have the statistical power to properly estimate these measures and their change across the two-year period. The matched control groups also provide the benchmark for the expected effect size for the growth in this neurodevelopmental factor within their respective socioeconomic regimes. Therefore, the lack of significant increase in the neurodevelopmental factor within the high-pollution groups in our study is not due to low statistical power. The fact that it is significantly smaller compared to the control groups supports the hypothesis that neurocognitive development is less than expected in the high-pollution groups.

The same neural indices are often related to multiple behaviors [Genon et al., 2018]. Thus, studies that assess only neural indices as a function of pollution or any other environmental factors absent any observed behavioral performance or self-report measures may suffer from reverse inference problem when discussing their data-driven neural findings. Therefore, another strength of the current study is including longitudinal N-back and NIH-Toolbox cognitive task performance data in the analyses. The primary PLS latent variables in both PM2.5 and O3 studies loaded strongly on both neurodevelopmental and task performance variables. This covariation within the same LV (rather than separate orthogonal LVs loading on neural vs. task performance variables) suggests that the neural and behavioral variations are related (adjusted for all covariates). Notably, however, covariation within the same LV does not distinguish the directionality of relationships within the neurocognitive factor (i.e., neural → performance, performance → neural, or both being consequences of an underlying biobehavioral factor).

The associated neurocognitive development latent variable that separated the high-pollution vs. matched low-pollution control groups across the two years was similar between the two analyses, despite their dissimilar demographic compositions (groups had lower SES than total sample in the PM2.5 analysis but higher SES than total sample in the O3 analysis). Additionally, the results were unchanged after adjusting for neighborhood economic disadvantage (area deprivation index [Kind et al., 2018]). This suggests that the interruption to the neurocognitive maturation process from pre- to early adolescence associated with these pollutants is potentially different from that associated with low family or neighborhood resources [Kind et al., 2018]. This underscores the value of investigating associations between neurocognitive measures and specific environmental exposures [Berman et al., 2019; Hyde et al., 2020] in addition to aggregated measures of co-occurring experiences and environments [Keller et al., 2024].

There are at least three limitations to the current study. First, the measures of air pollution are based on concentrations in ambient air (1Km^2^ around the place of residence at baseline) and do not fully reflect the participant’s actual exposure to the pollutants. Second, pollutants like PM2.5 include many components and could vary across regions in terms composition, thus further breakdown of these components should be considered in future work when studying their potential neurotoxicant effects. Finally, in order to tackle the propensity differences between high-PM2.5 and low-PM2.5 groups of youth in the sample, we had to exclude the high-PM2.5 participants that were not represented among the low-pollution sub-sample. Future studies aiming to isolate the PM2.5 air pollution impacts in a more representative way than the current study can recruit more youths of color and with extremely low family income and parental education who reside in the low-PM2.5 areas in the US compared to the ABCD Study.

In conclusion, this study isolated the associations of high PM2.5 and high O3 exposures with multimodal indices of neurocognitive development from pre- to early-adolescence. Using whole-cortex neurodevelopmental as well as cognitive performance measures, we found that normative neurocognitive maturation in this period is attenuated among youth residing in areas with high levels of PM2.5 and O3 air pollution compared to their peers. As healthy cognitive development is central to multiple domains of resilience [Masten & Cicchetti, 2016] and academic achievement [Peng et al., 2020], more effort should be directed toward reduction of these pollutants in residential and school areas.

## Acknowledgements

O.K. was supported by the National Institute on Drug Abuse K01 DA059598. A.W. was supported by National Institute on Drug Abuse K01 DA051561.

## ABCD study acknowledgements

Data used in the preparation of this article were obtained from the Adolescent Brain Cognitive DevelopmentSM (ABCD) Study (https://abcdstudy.org) Release 5.1, held in the NIMH Data Archive (NDA). This is a multisite, longitudinal study designed to recruit more than 10,000 children age 9-10 and follow them over 10 years into early adulthood. The ABCD Study® is supported by the National Institutes of Health and additional federal partners under award numbers U01DA041048, U01DA050989, U01DA051016, U01DA041022, U01DA051018, U01DA051037, U01DA050987, U01DA041174, U01DA041106, U01DA041117, U01DA041028, U01DA041134, U01DA050988, U01DA051039, U01DA041156, U01DA041025, U01DA041120, U01DA051038, U01DA041148, U01DA041093, U01DA041089, U24DA041123, U24DA041147.

Additional support for linked external data used in this work was made possible from NIEHS R01-ES032295 and R01-ES031074.

## Code and data availability

Scripts are available at https://github.com/okardan/ABCD_Air. Data are available from the Adolescent Brain Cognitve Development Study after approval of data use.

## Conflicts of interest

No conflicts to report.

## Notes

### Competing Interest Statement

The authors have declared no competing interest.

### Summary of Updates

1) The analytic approach has been changed from multiple regressions with continuous pollution to partial least squares with high-pollution vs. matched control groups over time. 2) The air pollution measures have been updated as per ABCD Study’s update of the measures from Release 2.0 (which used 10x10Km grid) to the later Releases where a 1x1Km grid was used instead around the participant’s residence. 3) Structural and functional cortical neurodevelopmental measures are now used instead of the cognitive composite network predictors. 4) Pollutants (PM2.5, NO2, and O3) are now separately assessed.

